# Induce Pluripotency via Specific distal enhancer-promoter associations

**DOI:** 10.1101/2021.06.09.447809

**Authors:** Xiusheng Zhu, Lei Huang, Dongwei Li, Jing Luo, Qitong Huang, Xueyan Wang, Yubo Zhang

## Abstract

Induced pluripotent stem cell (iPSC) technology promises to be an inexhaustible source of any type of cell needed for therapeutic and research purposes. It is unclear that how distal enhancer-promoter associations/ 3D chromatin conformation involving in the capacity of self-renewal and pluripotency maintenance. In this study, we have selected a few defined enhancer-promoter associations. After screening of enhancer specificity and activity individually, we design the different combinations and transfect these enhancers into the MEF cells. We simultaneously transfect 7 determined enhancers which represents various specific distal chromatin associations into a GFP tracing MEF cell line. We observe that the MEF cells start generating iPS-like clones at day 22. Importantly, our validations with three germ layer marker genes and *in vitro* experiments have further confirmed the pluripotency of these clones. Here, our study proposes a potential *de novo* method of a low-genetic risk iPS generation by introducing spatiotemporal distal chromatin associations. This result also paves out the way on utilizing 3D genomic information to alter cell identity and reprogramming for potential therapeutic strategy.

## Introduction

Induced Pluripotent Stem Cells (iPS) is a promising technology and derived from adult cells that have been reprogrammed back into an embryonic-like pluripotent state and it enables the development of an unlimited source of any type of cell needed for therapeutic purposes and other applications. It opens up unprecedented opportunities in regenerative medicine, disease modelling and drug discovery. Therefore, researchers have rapidly improved the techniques to generate iPSCs and lots of different methods have been developed since then. There are still a few shortcomings to limit its application, such as production efficiency, safety, carcinogenicity and etc^1–8^. Among them, one big concern is the permanent effects by introducing the endogenous gene/TFs would disturb the transcriptional factory in the cells. This might have some unpredicted consequences and inscrutable effects in the long run or next generation individual/cells^9^.

Three-dimensional (3D) genomics is emerging as a powerful tool to study the 3D structure and transcriptional regulation of eukaryotic genomes during the last decade^10^. Scientists have started realizing the interplay between transcription and genome conformation is a driving force for cell-fate decisions^11^. Through 3D genomics, we can bring the role of genome conformation in transcriptional regulation to the fore and understand the basic structure of DNA, the growth and development of organisms and the occurrence of diseases. However, at present, the researches still remain on describing the physical interactions/associations within various cell/organs. It is a major challenge to set up the solid connection between defined 3D genomic information and cell-fate decisions.

In our previous study, we have identified the physical transcriptional network existing in mouse embryonic stem cells (mESCs)^12^. The similar result is also described in mouse neural stem cells (mNSCs). So we recognize that the OSKM or Yamanaka factors (Oct4, Sox2, Klf4, and cMyc) are all connected via different chromatin associations. We find these enhancer-promoter associations show strong specificity comparing with other cells which is alike in OSKM’s gene expression patterns. Additionally, Nanog has also been reported to be essential for the establishment of both embryonic stem cells (ESCs) from blastocysts and iPSCs from somatic cells^13^, which is also involved in the network via specific interactions. In this study, we have selected these defined mES specific associations to explore the relationship between cell-fate decision and these specific chromatin associations. And we choose the classical iPSs technical mode as our research roadmap.

## Results

### Activation of Endogenous Pluripotent Gene

*Oct4*, *Sox2*, *Klf4* and *Nanog* were selected as targets because of their central roles in pluripotency maintenance and induction. In previous study, we have identified the specific chromatin associations of them^12^. Further, we verify their activities by considering other factors such as H3K27ac and H3K4me1 modification in pluripotent stem cells, and their potential to form promoter-enhancer loops mediated by Mediator complex^14–16^ (Fig.1A). Finally, we choose 9 candidate enhancers, namely OE1 (chr17:35622426-35623338), OE2 (chr17:35639453-35641591) (Oct4 enhancers); KE1 (chr4:55527598-55528553), KE2 (chr4:55663709-55664620) (Klf4 enhancer); SE1 (chr3:34656590-34657598), SE2 (chr3:34342406-34344100), SE3 (chr3:34562477-34563357) (Sox2 enhancer); NE1 (chr6:122652070-122654308), NE2 (chr6:122619081-122620092) (Nanog enhancers) as candidates for the following experiments.

**Figure.1.**
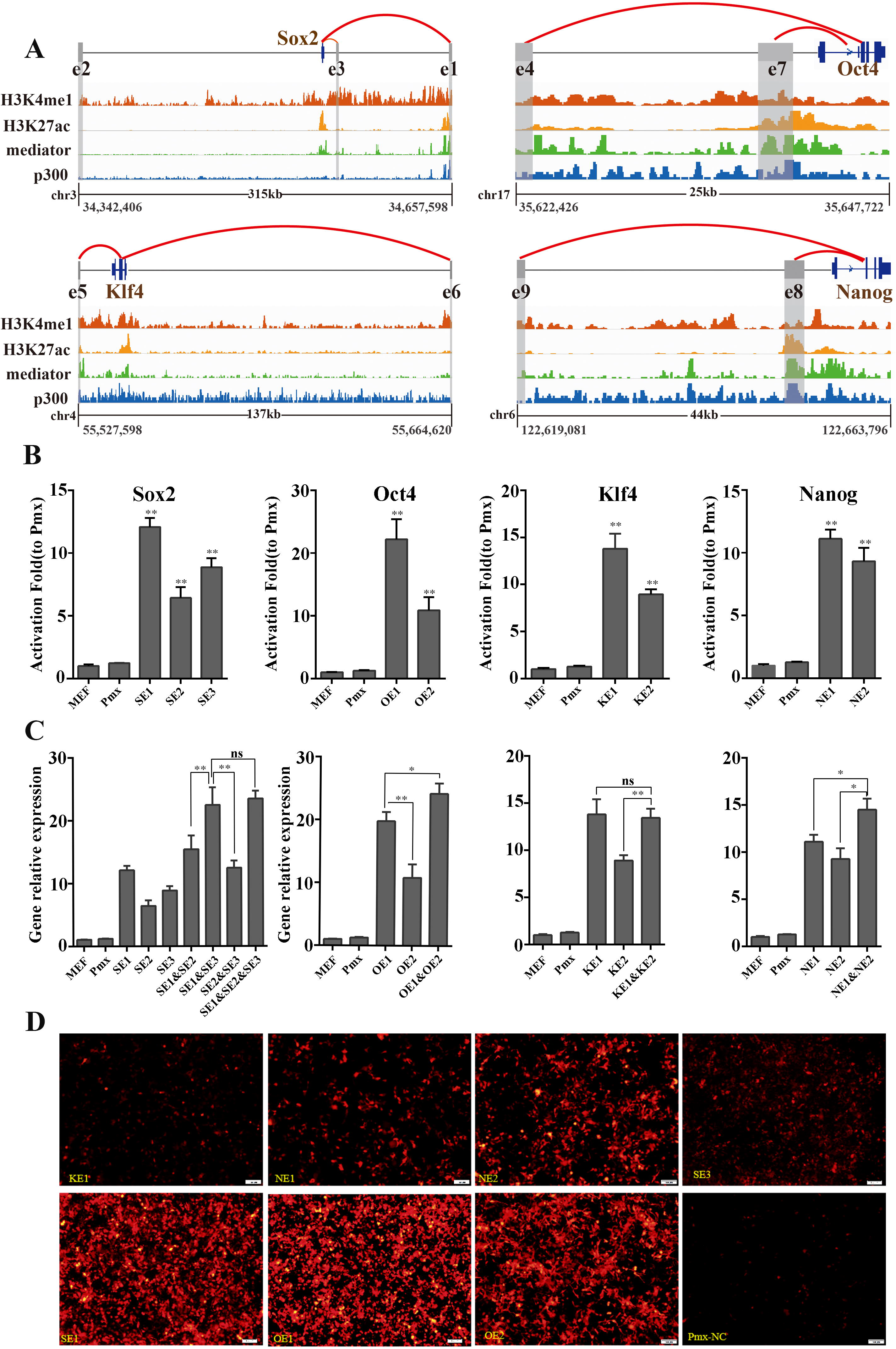
Gene-specific interaction enhancers activate endogenous gene expression. A: Describes the specific interaction between Oct4, Klf4, Sox2, Nanog enhancers and target genes, as well as the overlap of histone modifications, P300, and Mediator complex. B: The activation of exogenous specific enhancers on the transcription level of pluripotent genes. *: p<0.05; **: p<0.01. C. The effect of combined activation of different enhancer combinations on the transcription level of target genes. *: p<0.05; **: p<0.01, ns: There is no significant difference.

First, in order to test their transcriptional activities, we individually transfect these enhancers into the MEFs through a retroviral system^17^. We apply a transgenic Oct4 promoter driving GFP expression MEFs, OG2 MEF cells, as our model system. After 96 hours, we use qPCR to detect the expression level changes of these endogenous target genes. As expected, most of them can activate/enhance the expression of endogenous target genes at various levels (Fig. 1B). To reduce virus damage to cells and obtain higher activation efficiency, we further test the gene expression level of different enhancer combinations of individual pluripotent gene. The results show that the two enhancer combinations of *Oct4* gene have significantly higher transcriptional activation levels than individual enhancer. For *Sox2* gene, the combined activation of transcription levels of three enhancers is significantly higher than that of a single enhancer and SE1&SE2, SE2&SE3 combination. But there is no significant difference with the SE1&SE3. For the *Klf4* gene, the transcriptional activation level of the two enhancer combinations for the target gene is significantly higher than that of KE6, but there is no significant difference to KE5. The combination of two enhancers of *Nanog* gene has significantly higher transcriptional activation levels for target genes than individual one (Fig. 1C). Therefore, we choose OE1, OE2, KE1, SE1, SE3, NE1, NE2 as the candidates and use them to induce MEFs and to determine the relationship between the pluripotency and chromatin associations.

To double check its activity of these 7 chose enhancers, we construct them on a luciferase vector which has been widely used as a golden standard for this kind of validation experiments. The results verify the activity, and their expression levels are all significantly higher than the control group (Fig. 1D). These results are consistent with previous ones in zebrafish *in vivo*(ref). These indicate that exogenous enhancers can enhancer/activate transcription of the endogenous target genes.

### Induce Pluripotency via Specific distal interactions

We next try to determine whether pluripotency can be fully reactivated and established in MEFs following classical iPSs procedure. As mentioned before, in order to detect the phenomena more conveniently, we use OG2 MEF cells for the whole process. With OG2 MEF cells, when the endogenous *Oct4* is activated, the cells show a strong EGFP signal and can easily capture the signal via microscopes^18^.

To determine whether these specific distal interactions are sufficient to induce pluripotency with OG2 MEFs, we follow the efficient retroviral transfection iPSs protocol^17^ to integrate these enhancers into the MEFs. 48 hours after transfection, the MEFs medium has been replaced with iCD1 medium, which is recorded as day 0. Then we daily track and observe the morphological changes of these cells. In the study, reprogramming clusters start appearing on the 11th day (Fig. 2A). On 18th day, the emergence of clones can be observed (Fig. 2B). In order to determine more details regarding to the changes of Oct4 and Sox2 gene during the reprogramming process, we monitor their expression levels at different time points. The results showed that Oct4 gene expression has been constantly increasing during the reprogramming process. But, for Sox2 gene, it is lower on the 18th day than on the 11th day (Fig. 2C), which indicates the expression pattern of them is a little different during the process. The number of clones increased significantly on the 22nd day. To determine its pluripotency, we determine the signals of the EGFP expression under a fluorescence microscope, and the results present that they are all EGFP-positive ones (Fig. 2D). These data indicates that the pluripotency has been induced.

**Figure 2.**
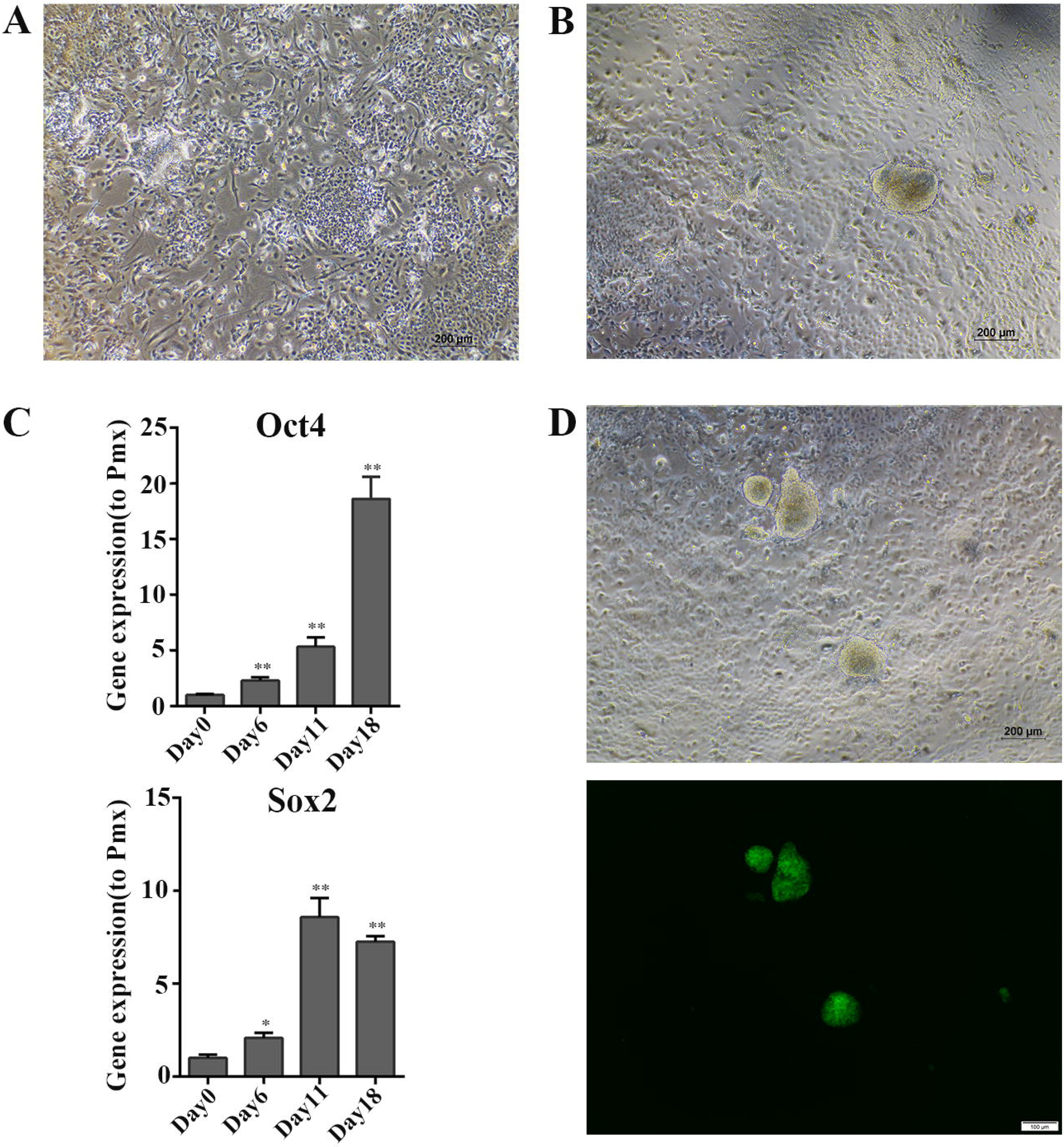
The specific interaction enhancer induces pluripotency network in MEFs. A. The MEFs induced on the 11th day showed cell aggregation. B. Clone-like cells appeared on the 18th day. C. The expression trend of the pluripotent genes Oct4 and Sox2 at different time points. D. The expression level of EGFP in clone-like cells observed by fluorescence microscope.

Next, in order to further confirm their differentiation ability *in vitro*, we pick some clones and culture them in suspension in a petri dish for 7 days. After the suspension culture process, these clones can form embryoid bodies (EBs) (Fig. 3A). In order to test whether they can differentiate into three primitive layers (endoderm, ectoderm, mesoderm), We continue to adhere to these EBs. In standard protocols, immunofluorescence experiments can be used to detect the specific expression pattern of three germ layers. Here, marker genes *SOX17* (endoderm), *β-Tubulin* (ectoderm), and *SMA* (mesoderm) are applied to determine the ability (Fig. 3B). In the results, it confirms that these EBs show specific expression pattern and have the potential to differentiate into other types of cells. Together, these results indicate that these specific distal interactions can induce the MEFs into a pluripotent status and form the iPSs like clones. Further, these clones can differentiate into three primitive layers and have the potential to differentiate into various types of cells *in vitro*.

**Figure 3.**
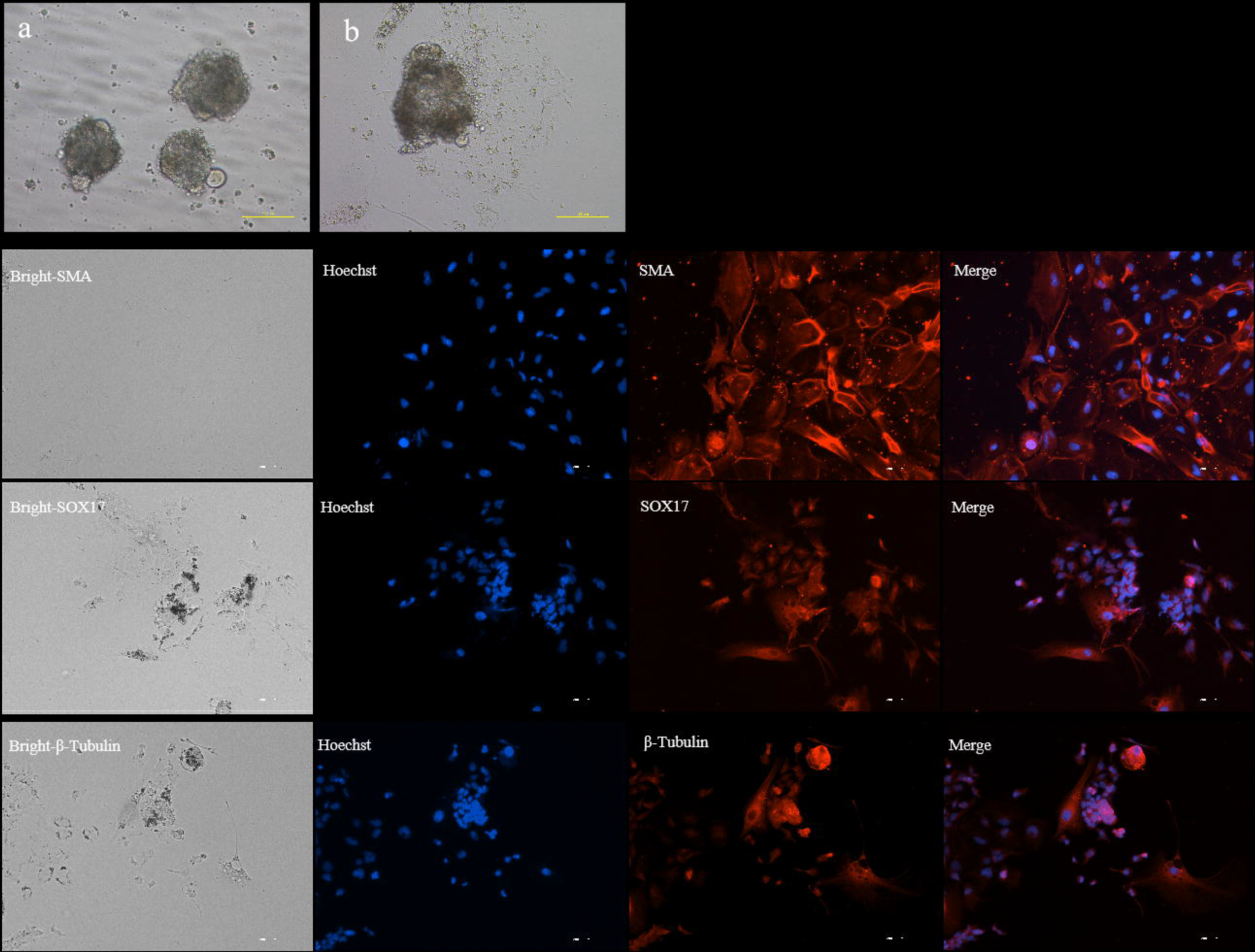
In vitro differentiation experiment of iPS-like cells. A. a. EBs formed by cloning-like cells in suspension culture for 7 days. b. EBs began to differentiate gradually to form many types of cells. B. Immunofluorescence experiments observed that EBs can differentiate spontaneously and express the marker genes of the three germ layers.

## Discussion

For putative regenerative applications, safety is always the foremost consideration for scientist. In traditional iPSs protocol, expression of OSKM factors is a key risk for sustained pluripotency, uncontrolled proliferation and tumorigenesis. It is known that TFs are coding genes and usually common to several cell types. They play a general role in gene regulation to determine the phenotypic characteristics of a cell^11^. This would bring unpredicted consequences and inscrutable effects and could limit its application. The distal enhancer-promoter chromatin associations present robust spatiotemporal specific in various studies^12,19,20^. The majority of enhancers are non-coding elements. Thus, manipulating them without disturb the gene/TF may dramatically reduce the risk and increase the safety. In this study, we choose the defined mESCs specific interactions of OSKM and follow the standard iPSs protocol. In the study, these specific interactions activate/enhance endogenous gene expression and result in iPS-like cells. Our method will lower the risk for iPSs application and provide an alternative route for iPSs generation.

3D chromatin architecture has been recognized in influencing gene regulation, cell fate decisions and evolution in the last decade. Also, it is known that it orchestrates transcriptional regulation act in a complex, multilayered and interconnected fashion. Different non-random spatial chromatin organization has been described based on various methodologies. However, all these studies are descriptive and hypothetical. Many of them imply or suspect a role for 3D genome conformation in the transcription factor-driven control of gene regulation and, consequently, cell-fate decisions. ChIA-PET is a unique technology which can bring along the function information of interested TFs with distal chromatin associations. In our previous study, we have identified physical connected transcription networks in mESCs and mNPCs. In current study, we apply these specific interactions to induce the iPSs following the standard protocol. Therefore, we have built a solid connection between 3D chromatin conformation and cell-fate decision here. We realize that cell identity can be determined by the spatiotemporal distal chromatin associations and a spatially organized genome. If that was the case, our method could propose some alternative therapeutic strategies by altering the cell-fate decision in cancer or other deadly diseases or ESCs/iPSs free route for regeneration therapies.

## Methods

### Cell culture

Plat-E cells purchased from ATCC, OG2 MEFs were from Dr. Baoming Qin’s laboratory, Plat E cells and MEFs were cultured in DMEM supplemented with 10% FBS and non-essential amino acid (NEAA).

All iPS-like cells and mouse ES cells were cultured in N2B27-2i/LIF medium(Neural Basal, F12,1%Glutamax, 1%NEAA, 1%Sodium pyruvate, 0.1mM β-me, 1%B27, 0.5%N2, 0.5% P/s(optional), mLif(1:8000), 3uM CHIR99021(1:4000), 1 uM PD0325901), MEFs are induced in iCD1 medium(Delicell,820250).

### Plasmid Construction

Pmx-mP-mCherry based on the modified pMX-GFP retroviral vector (Cell Biolabs, USA), is used in the study. Mini-promoter and mCherry (miniP-mCherry) fragments were obtained from modified pGL4.23 luciferase reporter vector (Promega, USA). Briefly, PGL4.23 was digested with XbaI/NcoI restriction enzymes to remove the original coding sequences of luc2 reporter gene and replace it with mCherry. The miniP-mCherry fragment was then amplified with primers containing NcoI or NotI restriction enzymes. Synthetic DNA fragment was cloned into NcoI and NotI digested pMX-GFP vector. The resulting screening vector was named pmx-mchery.Then double chop with Hind III and xhoI to get a linearized vector that can be attached to the input library.

### Retroviral Preparation and Transduction

Prepare Plat-E cells one day in advance, digest them with 0.25% pancreatin, and count them. Inoculate about 8×106/100mm plate of cells, wait until the cell density is about 80%-90%, transfect with lipo3000, change 10ml DMEM 10% FBS medium after 12 hours, and perform MEFs cell plating after 24 hours. The plating density was 5×104/well. After 12 hours, the supernatant of Plat-E cells was collected and filtered into a centrifuge tube with a 0.45μM filter membrane. After adding polybrene with a final concentration of 8μg/ml, the MEFs medium was replaced with fresh medium. Add the virus supernatant, add 1 ml of each enhancer virus supernatant (if it is a concentrated virus solution, reduce the corresponding multiple), Pmx-mCherry virus infection MEFs as a control group. A second virus transfection was carried out after 24 hours, and the medium was changed to iCD1 mouse induction medium after another 24 hours.

### Gene expression analysis

The cells were digested with 0.25% trypsin, and washed twice with PBS to collect the cells. The total RNA of the cells was extracted with the kit (Vazyme, RC112). Each reaction was reversed with 1 μg total RNA, and SYBR Green SuperMix (Vazyme, R323) on the 7500 Fast Real-Time PCR System (Applied Bio-systems). The pluripotent gene primer sequence is as follows: mOct4-F: ATGAGGCTACAGGGACACCTT,mOct4-R:GTGAAGTGGGGGCTTCCATA; mKlf-F:GTGCCCCGACTAACCGTTG, mKlf-R:GTCGTTGAACTCCTCGGTCT; mSox2-F:GCGGAGTGGAAACTTTTGTCC,mSox2-R:CGGGAAGCGTGTACTTA TCCTT;mNanog-F:AGGACAGGTTTCAGAAGCAGA,mNanog-R:CCATTGCTAG TCTTCAACCACTG;mGAPDH-F:AACTTTGGCATTGTGGAAGG,mGAPDH-R:A CACATTGGGGGTAGGAACA.

### In vitro differentiation experiment

Suspension culture of mouse ips-like cells in N2B27-2i/LIF medium for 7 days, and then continue to adhere to culture to differentiate into cells of different germ layers.

### Immunofluorescent Staining

Remove the cell culture medium, wash 3 times with PBS, and fix with 4% paraformaldehyde for 30 min at room temperature; wash 3 times with blocking solution (PBS containing 100 mmol/L glycine and 0.3% BSA), then add 1% Tritonx-100 Permeabilization at room temperature for 15 minutes; then add 5% bovine serum albumin (BSA) and block for 2 hours at room temperature; wash 3 times with TBP (Tritonx-BSA-PBS), add primary antibody and incubate overnight at 4 ℃; rewarm at room temperature the next day 20 min, wash with TBP 3 times, add secondary antibody and incubate for 1 h at room temperature; wash 3 times with TBP, then observe under a fluorescence microscope.

### Enhancer activity verification

The Pmx-mP-mCherry plasmid was double digested with endonucleases Hind III and XhoI, and the enhancer sequence to be verified was ligated to the vector by homologous recombination(Vazyme, Nanjing, China). Transfect the constructed plasmid into the cells(Invitrogen, Cat. No. L3000075), and observe the expression under a fluorescent microscope 24 hours later.

## Supporting information

enhancer site

## Acknowledgments

Thanks to the cell line provided by Baoming Qin Laboratory, Guangzhou Institute of Health, Chinese Academy of Sciences. Thanks to Dr. Ben Huang’s laboratory of Guangxi University for their help in the in vitro differentiation experiment. Thanks to Fei Fang for his help in cell culture experiments. Dr. Jun Yin reviews out manuscript.

## Author contributions

Y.Z., X.Z. and L.H. conduct and design the experiments. X.Z., D.L performed the experiments. X.W. help X.Z performed the experiments. Q.H, and J.L is responsible for photo editing and typesetting. The manuscript is written by X.Z. and Y.Z. All authors read and approved the final manuscript.

